# Asparagine signals mitochondrial respiration and can be targeted to impair tumour growth

**DOI:** 10.1101/2020.03.17.995670

**Authors:** Abigail S. Krall, Peter J. Mullen, Felicia Surjono, Milica Momcilovic, Ernst W. Schmid, Christopher J. Halbrook, Apisadaporn Thambundit, Steven D. Mittelman, Costas A. Lyssiotis, David B. Shackelford, Simon R.V. Knott, Heather R. Christofk

## Abstract

Mitochondrial respiration is critical for cell proliferation. In addition to producing ATP via the electron transport chain (ETC), respiration is required for the generation of TCA cycle-derived biosynthetic precursors, such as aspartate, an essential substrate for nucleotide synthesis. Because mTORC1 coordinates availability of biosynthetic precursors with anabolic metabolism, including nucleotide synthesis, a link between respiration and mTORC1 is fitting. Here we show that in addition to depleting intracellular aspartate, ETC inhibition depletes aspartate-derived asparagine and impairs mTORC1 activity. Providing exogenous asparagine restores mTORC1 activity, nucleotide synthesis, and proliferation in the context of ETC inhibition without restoring intracellular aspartate in a panel of cancer cell lines. As a therapeutic strategy, the combination of ETC inhibitor metformin, which limits tumour asparagine synthesis, and either asparaginase or dietary asparagine restriction, which limit tumour asparagine consumption, effectively impairs tumour growth in several mouse models of cancer. Because environmental asparagine is sufficient to restore proliferation with respiration impairment, both *in vitro* and *in vivo*, our findings suggest that asparagine synthesis is a fundamental purpose of mitochondrial respiration. Moreover, the results suggest that asparagine signals active respiration to mTORC1 to communicate biosynthetic precursor sufficiency and promote anabolism.

## Introduction

Recent literature has demonstrated that asparagine is important for amino acid homeostasis, maintenance of mTORC1 activity, and tumour progression^1–3^. Asparagine regulation of mTORC1 activity and downstream anabolism may explain the clinical efficacy of extracellular-acting asparaginase in the treatment of leukaemias that obtain most of their asparagine from the environment. Most solid tumours, however, are capable of synthesizing asparagine via asparagine synthetase (ASNS), making them less responsive to asparaginase^4^. In addition, elevated ASNS expression, and presumably increased *de novo* asparagine synthesis, accompanies asparaginase resistance in leukaemia^5^. Because cells can obtain asparagine from the environment or synthesize asparagine *de novo*, targeting both sources may be required to effectively exploit tumour asparagine dependence.

Cellular respiration couples nutrient oxidation to ATP production through oxidative phosphorylation. Although most cancer cells convert the majority of consumed glucose to lactate, concurrent respiration is essential: suppressing respiration through ETC inhibition impairs proliferation^6–9^. Recent literature has shown that ATP synthesis via the ETC is dispensable for cancer cell proliferation. Rather, aspartate synthesis requirements explain the reliance of proliferating cells on respiration^6, 7^. Electron acceptors are limiting upon ETC inhibition, resulting in compromised NAD+ recycling, impaired flux through the TCA cycle, and depletion of TCA cycle-derived aspartate. Supplementing cell culture medium with aspartate rescues the proliferation impairment caused by treatment with various ETC inhibitors without rescuing the redox state^6, 7^. These studies concluded that proliferating cells require respiration primarily for aspartate synthesis and aspartate-dependent nucleotide synthesis.

However, in addition to contributing carbon and nitrogen to nucleotide synthesis, another cellular use of aspartate is as a substrate for ASNS, which converts aspartate and glutamine to asparagine and glutamate (Figure 1a). This raises the possibility that aspartate synthesis in proliferating cells may also have another benefit: maintenance of cellular asparagine levels. Here we show that ETC inhibition is an effective means of impairing *de novo* asparagine synthesis. Moreover, we find that exogenous asparagine rescues proliferation impairment with ETC inhibition without rescuing intracellular aspartate levels, suggesting that aspartate rescue of proliferation upon ETC inhibition may be due, at least in part, to its conversion to asparagine via ASNS. Finally, using various mouse models of cancer, we provide evidence that combining ETC inhibition with treatments that limit environmental asparagine availability, such as asparaginase or dietary asparagine restriction, may be effective as a cancer therapy.

**Figure 1.**
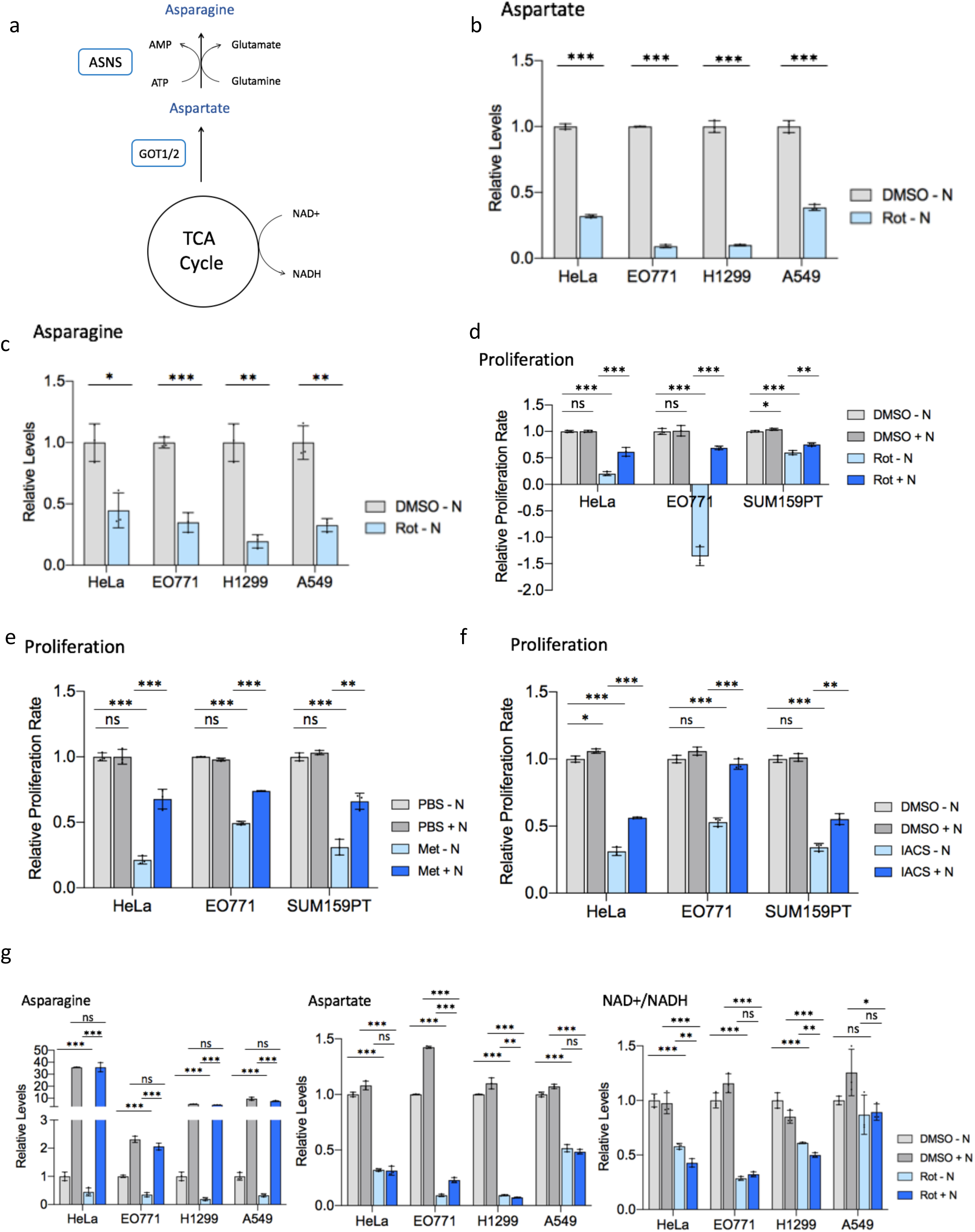
Asparagine restores proliferation in the context of ETC inhibition. **a**, Schematic diagramming asparagine synthesis from TCA cycle-derived aspartate. **b**, Relative levels of intracellular aspartate 6 hours post-treatment with rotenone (Rot) or vehicle (DMSO) in asparagine-free medium. **c**, Relative levels of intracellular asparagine 6 hours post-treatment with rotenone or DMSO in asparagine-free medium. **d**, Relative proliferation rate of indicated cell lines with rotenone or DMSO treatment in the presence or absence of 0.1 mM exogenous asparagine (N). **e**, Relative proliferation rate of indicated cell lines with metformin (Met) or vehicle control (PBS) treatment in the presence or absence of 0.1 mM exogenous asparagine (N). **f**, Relative proliferation rate of indicated cell lines with IACS-010759 (IACS) or vehicle control (DMSO) treatment in the presence or absence of 0.1 mM exogenous asparagine (N). **g**, Relative levels of intracellular asparagine, aspartate, and relative NAD+/NADH ratio in the indicated cell line 6 hours post-treatment with rotenone or DMSO in the presence or absence of 0.1 mM exogenous asparagine (N). Data are mean +/- s.d. (n = 3 independent experiments). P value determined by unpaired two-tailed t-test: *p<0.05; **p<0.01; ***p<0.001; ns, not significant.

### Asparagine rescues proliferation with ETC inhibition

ETC inhibition has been shown to reduce intracellular aspartate levels, with exogenous aspartate supplementation to cell culture medium rescuing ETC inhibition-mediated proliferation impairment^6, 7^. Aspartate is a precursor for protein, nucleotide, and asparagine synthesis. The extent to which each of these biosynthetic contributions is limiting for proliferation upon cellular aspartate depletion with ETC inhibition is unclear. Because aspartate is a direct substrate for the ASNS reaction (Fig. 1a), we examined the impact of ETC inhibition on intracellular asparagine. We confirmed that ETC inhibition with complex I inhibitor rotenone reduces intracellular aspartate levels in a panel of cancer cell lines (Fig. 1b). However, in addition to depleting aspartate, inhibition of the ETC also reduces intracellular asparagine (Fig. 1c).

To determine whether intracellular asparagine depletion contributes to the proliferation impairment observed with ETC inhibition, we inhibited the ETC in the presence or absence of exogenous asparagine supplementation in a panel of cancer cell lines. Physiologic levels of exogenous asparagine (100 uM) rescue the proliferation defect caused by ETC complex I inhibitors rotenone (Fig. 1d, Supplementary Fig. 1a), metformin (Fig. 1e, Supplementary Fig. 1b), and IACS-010759 (Fig. 1f) in most examined cell lines cultured in pyruvate-free DMEM, to a degree comparable to supraphysiologic levels of exogenous aspartate (20 mM)^6, 7^ (Supplementary Fig. 1c-d). Asparagine similarly rescues proliferation with inhibition of ETC complex III (antimycin A) and complex V (oligomycin) (Supplementary Fig. 1e). A minority of cell lines, including H1299 and A549 lung cancer cells, do not exhibit proliferative rescue by asparagine in standard DMEM (Supplementary Fig. 2a-b); however, rescue by asparagine is observed in these cell lines when DMEM is supplemented with physiologic concentrations of additional non-essential amino acids (see Supplementary Table I), including proline and taurine, which are depleted specifically in H1299 and A549 with rotenone treatment (Supplementary Fig. 2c-e). These results suggest amino acids in addition to asparagine and aspartate may be limiting for growth with ETC inhibition in certain cell culture conditions, and cell environment may therefore affect the extent to which asparagine restores proliferation. Importantly, exogenous aspartate supplementation rescues intracellular asparagine levels in the context of ETC inhibition (Supplementary Fig. 3a); however, exogenous asparagine rescues ETC inhibition proliferation defects without rescuing intracellular aspartate levels or the NAD+:NADH ratio (Fig. 1g, Supplementary Fig. 3a). Moreover, the extent of proliferation rescue by exogenous asparagine and aspartate is not additive (Supplementary Fig. 3b), suggesting that aspartate and asparagine may rescue proliferation through a common mechanism. Taken together, these data suggest that 1) asparagine, not aspartate, may be limiting for proliferation upon ETC inhibition; 2) aspartate rescue of proliferation upon ETC inhibition may be due to conversion of aspartate to asparagine via ASNS; and 3) asparagine synthesis may be a fundamental purpose of mitochondrial respiration in proliferating cells.

### Asparagine communicates mitochondrial respiration to ATF4 and mTORC1

We next examined the mechanism by which asparagine rescues proliferation with ETC impairment. Intracellular asparagine levels influence the activities of activating transcription factor 4 (ATF4) and mTOR complex I (mTORC1)^1^, both of which can impact cell proliferation. ATF4 plays a vital role in cellular homeostasis, activating transcription of genes involved in amino acid transport and synthesis when intracellular amino acids are depleted^10^. In particular, ASNS is a well-documented target gene of ATF4, and ATF4 is activated by asparagine depletion in order to increase *de novo* asparagine synthesis via ASNS^1, 10^. Chronic ATF4 activation, however, promotes apoptosis. mTORC1 activity positively regulates cell proliferation by activating anabolic processes required for cell division. We previously showed that intracellular asparagine depletion impairs mTORC1 activity and downstream anabolic processes, such as nucleotide synthesis^1^.

Because ETC inhibition results in asparagine depletion, we assessed ATF4 and mTORC1 activities with various ETC inhibitors in the presence or absence of exogenous asparagine. Complex I inhibition with rotenone or IACS-010759 increases ATF4 protein levels and decreases mTORC1 activity, as assessed by phosphorylation of mTOR target gene S6K, in a panel of cancer cell lines (Fig. 2a-b, Supplementary Fig. 4a). Exogenous asparagine supplementation rescues both ATF4 and mTORC1 activities in all examined cell lines. Moreover, abolishing the ability of the cell to synthesize asparagine via CRISPR-mediated ASNS knockout eliminates ATF4 and mTORC1 sensitivities to ETC activity and aspartate (Fig. 2c-d, Supplementary Fig. 4b-c). These results suggest that the altered ATF4 and mTORC1 activities observed with respiration impairment are primarily a response to impaired asparagine synthesis. It also raises the possibility that ATF4 activation and/or compromised anabolism downstream of mTORC1 contribute to the proliferation impairment observed with ETC inhibition in the absence of asparagine supplementation.

**Figure 2.**
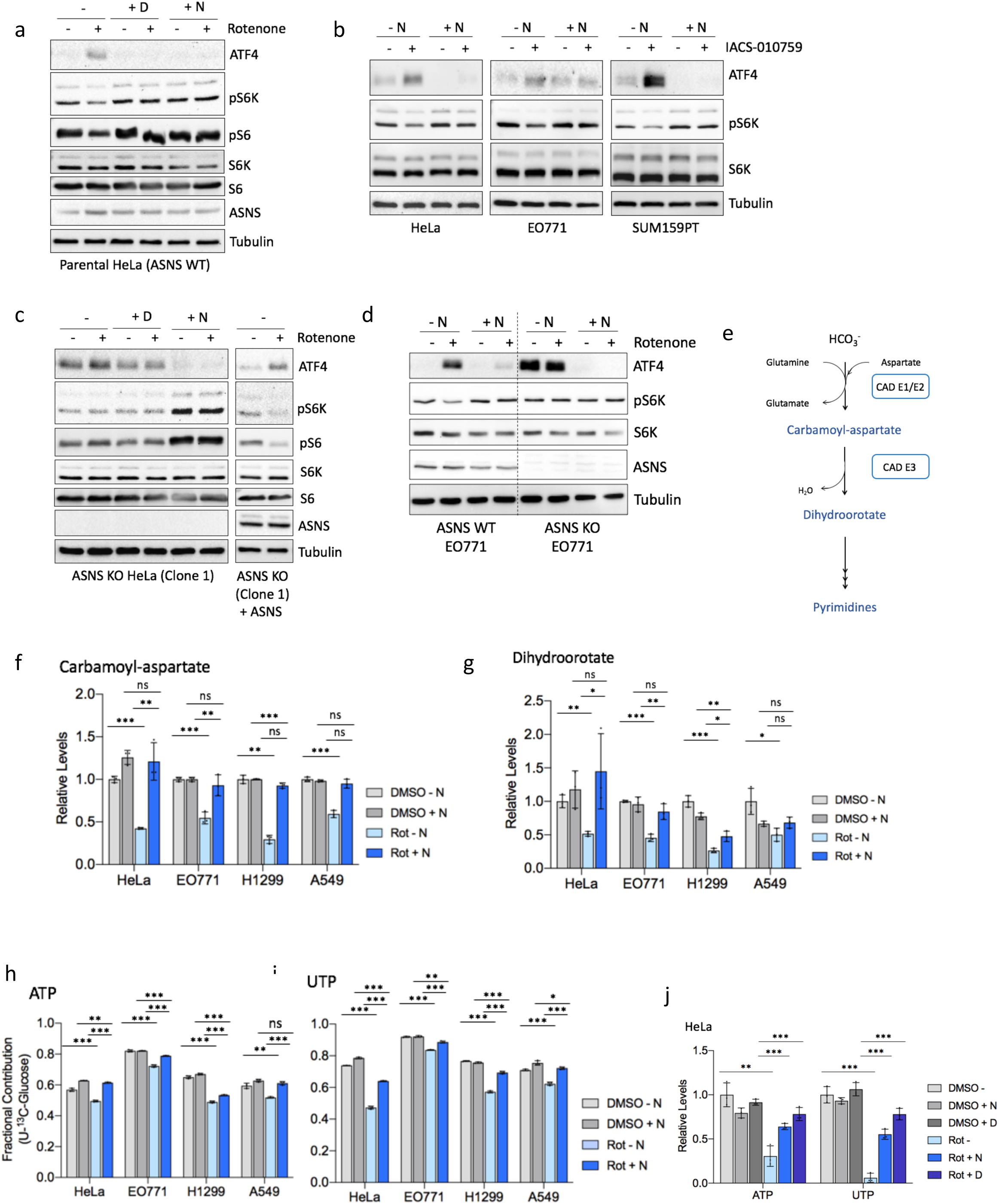
Asparagine relays mitochondrial respiration to ATF4 and mTORC1. **a**, Immunoblot of HeLa lysates 6 hours post-treatment with 50 nM rotenone (Rot) or DMSO in the presence or absence of 20 mM aspartate (D) or 0.1 mM asparagine (N). Lysates were immunoblotted for ATF4, mTORC1 activation markers phospho-Thr389 S6K and phospho-Ser235/6 S6, total S6K and S6, ASNS, and tubulin. **b**, Immunoblot of lysates after IACS-010759 (IACS) or DMSO treatment of HeLa (6 hours), E0771 (3 hours), and SUM159PT (6 hours) cells. **c**, Left, Immunoblot of HeLa ASNS KO (clone 1; see Supplementary Fig. 4b for ASNS KO clone 2) lysates 6 hours post-treatment with 50 nM rotenone or DMSO in the presence or absence of 20 mM aspartate (D) or 0.1 mM asparagine (N); Right, ASNS was restored in HeLa ASNS KO cells with CMV-driven ectopic expression. Immunoblot shows lysates 6 hours post-treatment with 50 nM rotenone or DMSO in unsupplemented DMEM. **d**, Immunoblot of E0771 WT or ASNS KO lysates 3 hours post-treatment with 50 nM rotenone or DMSO in the presence or absence of 0.1 mM asparagine (N). **e**, Schematic diagramming carbamoyl-phosphate synthetase 2 (CAD) activity in pyrimidine synthesis. Relative levels of CAD products, carbamoyl-aspartate (**f**) and dihydroorotate (**g**) in the indicated cell line 6 hours post-treatment with rotenone or DMSO presence or absence of 0.1 mM exogenous asparagine (N). Fractional contribution of U-^13^C-glucose to ATP (**h**) and UTP (**i**) in the indicated cell line 6 hours post-treatment with rotenone or DMSO presence or absence of 0.1 mM exogenous asparagine. Medium was replaced with DMEM containing 10 mM U-^13^C-glucose at the same time as rotenone treatment. **j**, Relative levels of intracellular ATP and UTP in HeLa cells 48 hours post-treatment with 50 nM rotenone or DMSO in the presence or absence of 0.1 mM asparagine (N) or 20 mM aspartate (D). Data are mean +/- s.d. (n = 3 independent experiments). P value determined by unpaired two-tailed t-test: *p<0.05; **p<0.01; ***p<0.001; ns, not significant.

It was previously reported that ETC inhibition causes nucleotide deficiency^6, 7^. Aspartate is a substrate for nucleotide synthesis, with aspartate atoms being incorporated into both purines and pyrimidines, and it has been suggested that nucleotide deficiency with ETC impairment is a result of reduced aspartate availability. However, purine and pyrimidine synthesis are also responsive to mTORC1 activity^11, 12^. For instance, mTORC1 promotes the activity of carbamoyl-phosphate synthase 2 (CAD), the rate-limiting enzyme of pyrimidine synthesis^11^ (Fig. 2e). Because we previously demonstrated that asparagine promotes mTORC1-mediated CAD phosphorylation to influence nucleotide levels^1^, we asked whether impaired nucleotide synthesis upon ETC inhibition may be a result of reduced mTORC1-mediated *de novo* nucleotide synthesis. Consistent with asparagine depletion, CAD phosphorylation on serine 1859 is reduced with rotenone treatment and rescued with exogenous asparagine in E0771 breast cancer cells (Supplementary Fig. 4d). Moreover, treatment with rotenone reduces levels of CAD products carbamoyl-aspartate and dihydroorotate (Fig. 2e-g), reduces glucose incorporation into purines and pyrimidines (Fig. 2h-i, Supplementary Fig. 4e-f), and reduces nucleotide levels in a panel of cell lines, in a manner that is rescued with exogenous asparagine (Fig. 2j). Although aspartate levels are reduced by ETC inhibition, and aspartate is a CAD substrate, asparagine rescues levels of CAD products and nucleotide synthesis without restoring intracellular aspartate (Fig. 1g). Furthermore, levels of glutamine, an additional CAD substrate, are not reduced with ETC inhibition (Supplementary Fig. 4g), suggesting that impaired nucleotide synthesis is unlikely to be caused by limited substrate availability. Taken together, these data suggest that with respiration impairment aspartate may be limiting for asparagine synthesis and asparagine-mediated mTORC1 signaling and anabolism, as opposed to being limiting for direct incorporation into nucleotides.

Although asparagine and aspartate both rescue HeLa proliferation at lower metformin doses (0.5 mM, a dose that impairs proliferation by 80%), aspartate, but not asparagine, rescues proliferation at a high metformin dose (2.5 mM) (Supplementary Fig. 4h). This suggests, that with enough ETC inhibition, aspartate does become depleted enough to be limiting for incorporation into nucleotides or protein, given that asparagine and asparagine-mediated signaling are insufficient to substitute for aspartate. This implies that the degree of aspartate depletion required to impair asparagine synthesis is lower than that required for aspartate to be limiting as a substrate for other aspartate-dependent processes, and that ASNS-dependent asparagine synthesis is a primary purpose of respiration-derived aspartate.

### The combination of metformin and asparaginase impairs tumour growth

The fact that cancer cell proliferation requires intracellular asparagine maintenance suggests that targeting this dependence may be an effective therapeutic strategy. Because tumours usually have the capacity for *de novo* asparagine synthesis, asparaginase, which hydrolyzes extracellular asparagine, only works to limit cancer cell proliferation in leukaemia with low ASNS activity. Simultaneously targeting both asparagine consumption and *de novo* asparagine synthesis may therefore be required to effectively exploit tumour asparagine dependence and impair tumour progression. In support of this approach, combining genetic silencing of ASNS with asparaginase-mediated depletion of blood asparagine has been shown to inhibit sarcoma growth *in vivo*^3^. Our *in vitro* results indicate that ETC inhibition is an effective non-genetic means to impair *de novo* asparagine synthesis. Complex I inhibitor metformin is a well-tolerated drug that is currently used in the clinic to treat patients with type 2 diabetes. We hypothesized that co-targeting cancer cell asparagine synthesis with metformin and asparagine consumption with asparaginase would reduce tumour growth.

To test this hypothesis, immunodeficient NOD scid gamma mice (NSG) mice harboring A549 lung subcutaneous xenografts were treated with a combination of metformin and asparaginase or with each drug individually. 5 IU/kg asparaginase effectively depleted circulating asparagine without affecting circulating glutamine (Supplementary Fig. 5a-b), and 250 mg/kg/day metformin resulted in serum metformin concentrations comparable to therapeutic levels in diabetes patients (Supplementary Fig. 5c)^13, 14^. While neither metformin nor asparaginase affects tumour growth independently, combining the two treatments significantly impairs A549 tumour growth (Fig. 3a-b, Supplementary Fig. 5d). Comparable results are observed with SUM159PT breast cancer xenografts in NSG mice (Fig. 3c). Given that these xenograft experiments were performed in immunodeficient mice, we verified that metformin combined with asparaginase does not affect an anti-tumour immune response. The combination does not impact anti-cancer activity of OT1 T cells *in vitro* (Supplementary Fig. 5e). Moreover, the combination of phenformin, a metformin-related biguanide that has been suggested to have anti-tumour activity in pancreatic cancer^15^, and asparaginase significantly reduces tumour growth in a syngeneic orthotopic model of pancreatic cancer in immune-competent mice (Fig. 3d). To determine whether the growth impairment with the metformin/asparaginase combination may be due to tumour asparagine depletion-mediated changes in signaling, mTORC1 activity was compared in A549 tumours across the treatment groups. Mice treated with the combination of metformin and asparaginase exhibit reduced tumour mTORC1 activity, as assessed by S6 phosphorylation on Ser235/236 and S6K electrophoretic mobility shift (Fig. 3e). These results indicate that extracellular asparagine availability via asparaginase and *de novo* asparagine synthesis via ETC inhibition impairs tumour growth.

**Figure 3.**
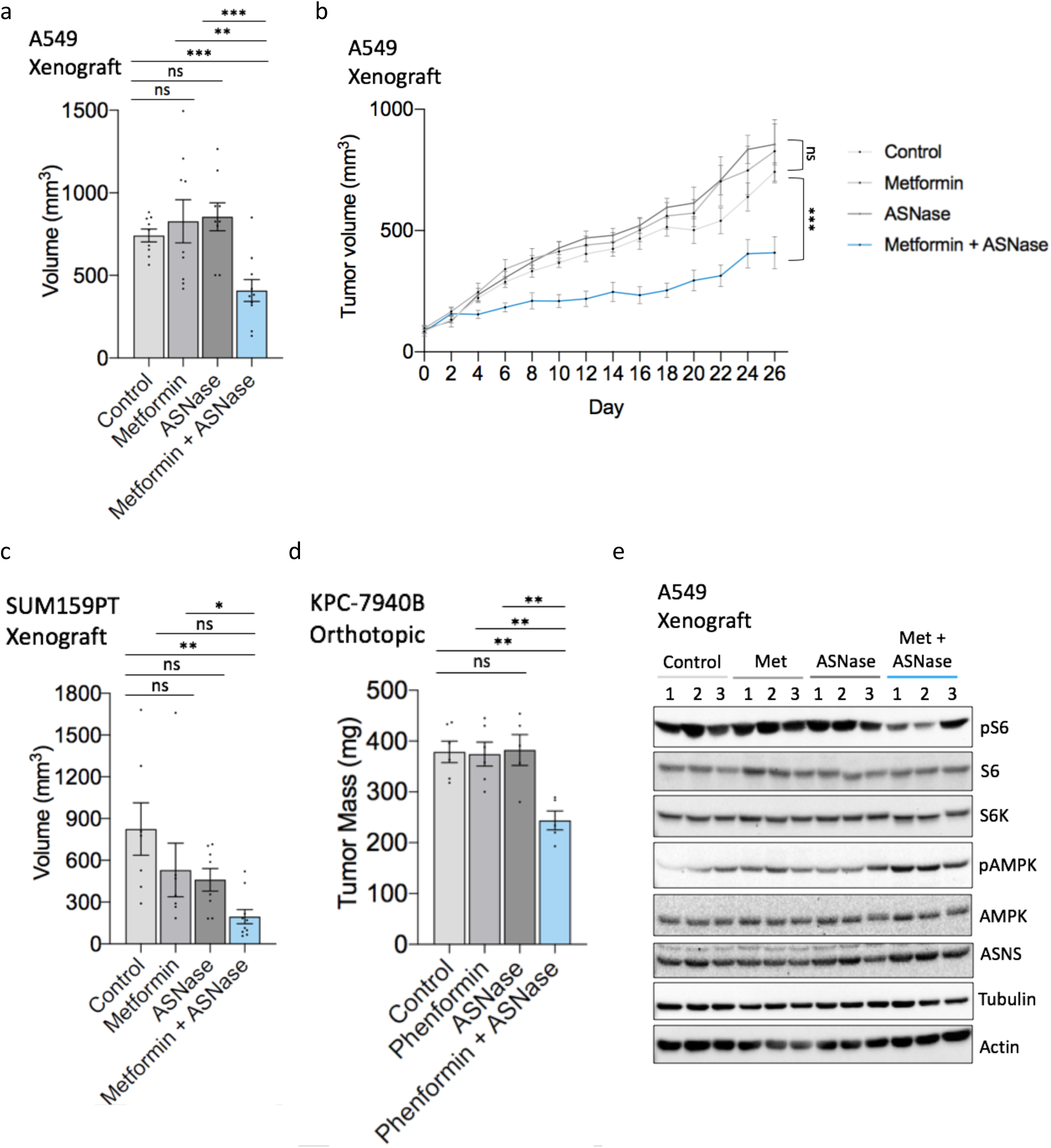
Combining metformin with asparaginase impairs tumour growth. **a**, Endpoint tumour volume (mm^3^) (day 26 of treatment) of A549 subcutaneous tumour xenografts in mice treated with metformin (250 mg/kg/day), asparaginase (ASNase) (5 IU/kg), the combination, or vehicle controls as determined by caliper measurements (n = 9-10). **b**, A549 tumour xenograft growth curves from metformin/asparaginase treatment start date through endpoint. **c**, Endpoint tumour volume (mm^3^) (day 21 of treatment) of SUM159PT subcutaneous tumour xenografts in mice treated with metformin (250 mg/kg/day), asparaginase (ASNase) (5 IU/kg), the combination, or vehicle controls as determined by caliper measurements. n = 7-10. **d**, Endpoint tumour mass of KPC-7940B orthotopic tumours treated with phenfomrin (1.7mg/mL), asparaginase (ASNase) (2 IU), the combination, or vehicle controls. n = 5-6. **e**, Immunoblot of lysates from metformin/asparaginase-treated A549 tumour xenografts shown in **a-b**. Lysates were immunoblotted for mTORC1 activation marker phospho-Ser235/6 S6, total S6K, total S6, phospho-Thr172 AMPK, total AMPK, ASNS, tubulin, and actin. The three middle-sized tumours of each treatment group were chosen as representatives. Data are mean +/- s.e.; P value determined by unpaired two-tailed t-test: *p<0.05; **p<0.01; ***p<0.001; ns, not significant.

### ETC inhibition combined with dietary asparagine restriction impairs tumour growth

It was recently shown that blood asparagine levels can be altered by the diet: mice fed chow with varying concentrations of asparagine exhibit dose-dependent differences in serum asparagine concentrations^2^. We therefore attempted modulating dietary asparagine as an alternative approach to asparaginase for limiting extracellular asparagine availability. We confirmed that feeding mice chow containing 0%, 0.6%, or 4% asparagine leads to corresponding differences in serum asparagine levels (Supplementary Fig. 6a). To determine whether a low-asparagine diet sensitizes tumours to metformin, NSG mice harboring A549 lung subcutaneous tumour xenografts were treated with or without metformin while fed a 0%, 0.6%, 4% asparagine diet. Although varying the amount of asparagine in the diet alone has no impact on tumour growth, metformin significantly impairs tumour growth in mice fed a 0% asparagine diet (Fig. 4a-b, Supplementary Fig. 6b-c). Importantly, the 0.6% and 4% asparagine diets rescue tumour growth in a dose-dependent manner, suggesting that metformin impairs tumour growth by limiting asparagine synthesis. A similar growth impairment was observed in A549 tumour xenografts when dietary asparagine restriction was combined with alternative complex I inhibitor IACS-010759 (Fig. 4c-d, Supplementary Fig. 6d), providing confidence that the phenotype we observed with metformin *in vivo* is due to metformin-induced inhibition of the ETC as opposed to other metformin targets. Moreover, similar to what is observed with the combination of metformin and asparaginase, dietary asparagine restriction combined with either metformin or IACS-010759 leads to a general decrease in tumour mTORC1 activity (reduced S6 phosphorylation on Ser235/236 and S6K phosphorylation on Thr389), and a signaling rescue is observed with dietary asparagine supplementation (Fig. 4e, Supplementary Fig. 6e-f).

**Figure 4.**
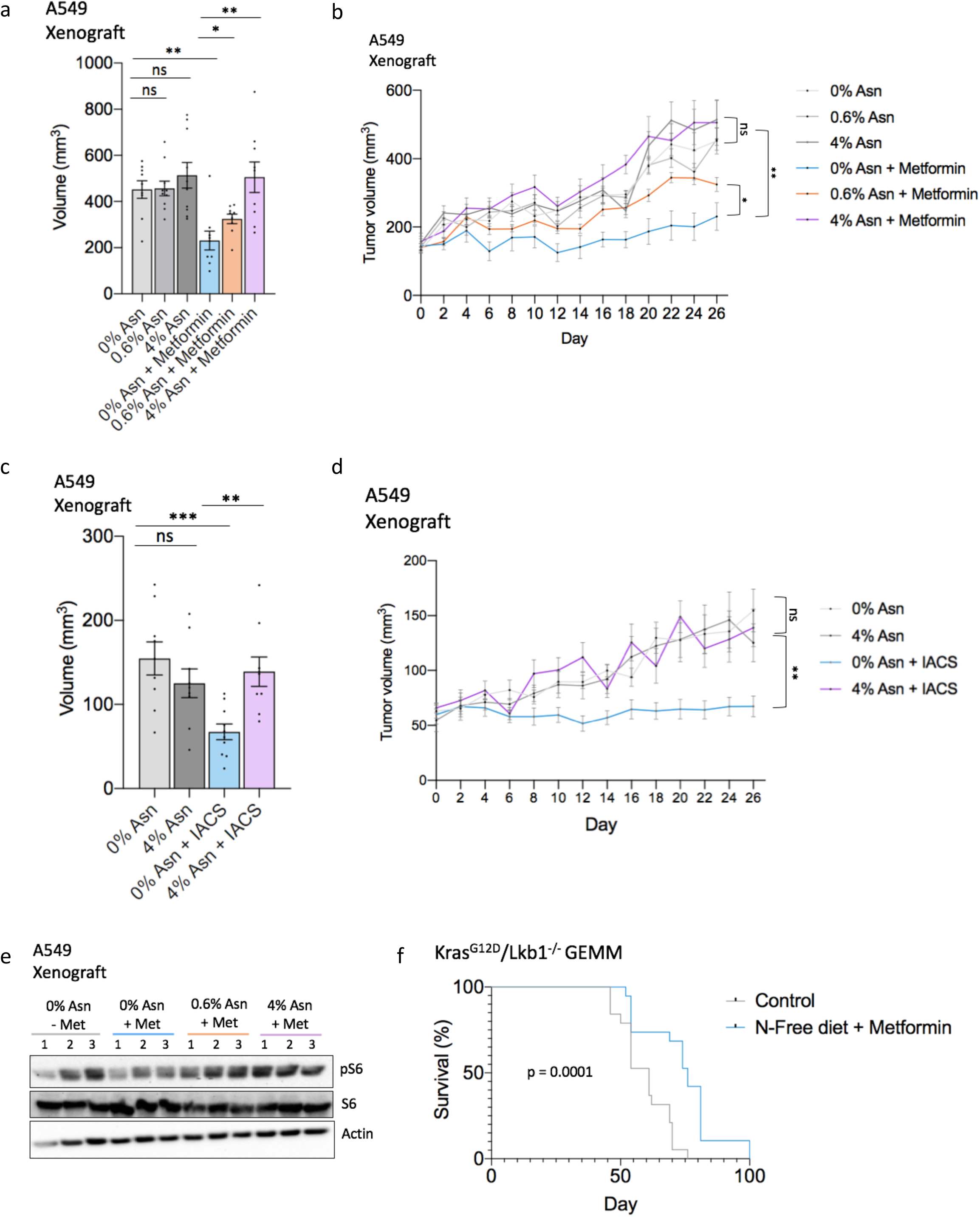
Combining metformin with dietary asparagine restriction impairs tumour growth. **a**, Endpoint tumour volume (mm^3^) (day 26 of treatment) of A549 subcutaneous tumour xenografts in mice treated with or without metformin and fed a diet containing 0%, 0.6%, or 4% asparagine, as determined by caliper measurements. n = 9-10. **b**, A549 tumour xenograft growth curve from metformin/asparagine diet start date through endpoint. **c**, Endpoint tumour volume (mm^3^) (day 26 of treatment) of A549 subcutaneous tumour xenografts in mice treated with IACS-010759 (IACS) (1 mg/kg) or vehicle and fed a 0% or 4% asparagine diet, as determined by caliper measurements. n = 9-10. **d**, A549 tumour xenograft growth curve from IACS-010759/asparagine diet start date through endpoint. **e**, Immunoblot of lysates from metformin/dietary asparagine restriction-treated A549 tumour xenografts shown in **a-b**. Lysates were immunoblotted for mTORC1 activation marker phospho-Ser235/6 S6, total S6, and actin. The three middle-sized tumours of the indicated treatment groups were chosen as representatives. **f**, Kaplan-Meier survival curve indicating percentage of mice with Kras^G12D^/Lkb1^-/-^-driven non-small cell lung cancer surviving over 100 days. Control mice were fed a 0.6% asparagine diet without metformin in the drinking water; the treatment group (N-free diet + metformin) was fed a 0% asparagine diet and treated with 250 mg/kg/day metformin in the drinking water. Treatment was initiated (day 0) two weeks after Cre-mediated tumour induction (see methods). n = 19; Error bars denote s.e. of the mean. For **a-d**, data are mean +/- s.e.; P value determined by unpaired two-tailed t-test. For **f**, P-value was calculated by Mantel-Cox test. *p<0.05; **p<0.01; ***p<0.001; ns, not significant.

Finally, we examined the efficacy of the combination of metformin plus dietary asparagine restriction in two immune-competent tumour models. Metformin combined with dietary asparagine restriction significantly extends survival in a Kras^G12D^/Lkb1^-/-^-driven genetically engineered mouse model (GEMM) of non-small cell lung cancer (Fig. 4f). However, the combination does not significantly impact tumour growth or survival in a syngeneic mouse model of breast cancer (E0771 cells) (Supplementary Fig. 7a-d). Interestingly, although the phenotype is not explained by our proposed mechanism, dietary asparagine restriction appears to synergize with PD1 inhibition and chemotherapy (cyclophosphamide) in this breast cancer model (Supplementary Fig. 7c-d). Taken together, our results suggest that combining ETC inhibition with treatments that reduce circulating asparagine (either asparaginase or dietary asparagine restriction) has *in vivo* efficacy and is a promising therapeutic strategy.

## Discussion

We provide evidence that asparagine synthesis is a fundamental purpose of mitochondrial respiration for proliferating cells. Two of the requisite substrates for ASNS production of asparagine – aspartate and ATP – are outputs of mitochondrial respiration. In most cell types examined, providing exogenous asparagine can substitute for ETC activity and aspartate to permit proliferation.

Because blood aspartate levels (0-15 uM)^16, 17^ do not allow adequate aspartate import in most cell types, aspartate availability and aspartate-derived nucleotide synthesis depend on mitochondrial respiration and TCA cycle flux. We and others^1, 18–20^ have shown that ETC impairment leads to inhibition of mTORC1 activity. Communication of mitochondrial respiration to mTORC1 may prevent mTORC1-mediated activation of anabolic processes, such as nucleotide synthesis, when TCA cycle-derived biosynthetic precursors, such as aspartate, are limiting. Impaired mitochondrial respiration has also been shown to activate ATF4^21–25^, and there is accumulating evidence that ATF4 promotes transcription of genes that support respiration and relief of mitochondrial stress^24, 25^. We show that exogenous asparagine is sufficient to rescue mTORC1 inhibition and ATF4 activation in the context of respiration impairment, which suggests that asparagine is a crucial cellular signal of mitochondrial respiration (Supplementary Fig. 8). Consistent with the idea that asparagine generation is a fundamental purpose of mitochondrial respiration, the fact that asparagine rescues ATF4 activity with ETC inhibition also suggests that ATF4 transcriptional regulation of genes involved in mitochondrial respiration and stress relief is for the purpose of generating asparagine precursors to restore asparagine synthesis.

We show that ETC inhibition synergizes with asparaginase or dietary asparagine restriction in several mouse models of cancer. This indicates that ETC inhibition effectively restricts asparagine synthesis both *in vitro* and *in vivo*. The data also present a promising therapeutic strategy that may be readily available for cancer patients. Metformin is a well-tolerated drug that is clinically approved for diabetes. Asparaginase has been used clinically to treat leukaemia for decades. The proposed strategy may allow for rapid translation of an effective combination therapy to the clinic.

Although the combination strategy was effective over the time frame of our experiments, we acknowledge the existence of potential resistance mechanisms. Cancer cells may develop resistance to the combination by increasing pyruvate and/or aspartate uptake. Pyruvate can restore redox homeostasis by acting as an electron acceptor to rescue proliferation upon ETC inhibition^6, 7, 26^, and therefore, increased pyruvate uptake could potentially restore intracellular aspartate and asparagine levels. Although supraphysiologic levels of pyruvate are needed to restore redox homeostasis *in vitro*, and blood aspartate levels are presumably insufficient to restore *de novo* asparagine synthesis, long-term monitoring of tumour growth with the combination treatment will be required to adequately assess risk and mechanism of resistance development.

## Acknowledgements

We thank Andrew Martinez, Barbara Nelson, and Galloway Thurston for assistance with the animal studies and all members of the Christofk lab for assistance and constructive feedback. C.J.H was supported by F32CA228328, P30DK034933, and K99CA241357. C.A.L. was supported by a 2017 AACR NextGen Grant for Transformative Cancer Research (17-20-01-LYSS); an ACS Research Scholar Grant (RSG-18-186-01); and 1R37CA237421. H.R.C. was supported by RO1 CA215185, RO1 AR070245, a research scholar grant (RSG-16-111-01-MPC) from the American Cancer Society, Jazz Pharmaceuticals VT# IST-16-10306, and the UCLA Jonsson Comprehensive Cancer Center and Eli and Edythe Broad Center for Regenerative Medicine Ablon Scholars Program, and the Most Promising Research Award from UCLA Health Innovation.

## Methods

### Cell lines and culture conditions

For standard passaging, EO771 cells (CH3 BioSystems) were cultured in RPMI supplemented with 10% FBS, 1% penicillin-streptomycin, 1% MEM NEAA (Gibco), and 10 mM HEPES; all other cell lines were cultured in DMEM supplemented with 10% fetal bovine serum and 1% penicillin-streptomycin. The LPS2 cell line was derived from a liposarcoma tumour sample^27^.

### ETC inhibition

ETC inhibition experiments were performed for all cell lines in pyruvate-free DMEM supplemented with 1% penicillin-streptomycin and dialyzed FBS (Life Technologies) unless otherwise indicated in Supplementary Table I. For experiments on H1299 and A549 cell lines, medium was supplemented with 400 uM proline, 500 uM alanine, 100 uM glutamate, and 150 uM taurine, unless otherwise stated.

### Proliferation assays

Cells were seeded in triplicate in six-well plates at 5×10^4^ cells per well 24 hours prior to experiment initiation. At the start of the experiment, triplicate wells were counted using a particle counter (day 0), and medium was replaced with 3 mL of pyruvate-free DMEM containing or lacking 0.1 mM asparagine or 20 mM aspartate in the presence of ETC inhibitor or vehicle. Final cell counts were obtained 2-4 days post-medium change, and the number of cell doublings post-day 0 was determined. Serum and ETC inhibitor information for each cell line can be found in Supplementary Table I.

### Intracellular metabolite extraction and analysis

Cells were seeded in six-well plates, and metabolites were extracted at 70–80% confluence. For glucose labeling experiments, medium was replaced for 6 hours with DMEM containing 10 mM U-^13^C-glucose (Cambridge Isotopes), 10 mM ^12^C-glutamine, and 10% dialyzed FBS in the presence or absence of rotenone. For H1299 and A549, medium was supplemented with proline, alanine, glutamate, and taurine (see Supplementary Table I for cell line-specific conditions). After two washes with ice-cold 150 mM ammonium acetate, pH 7.3, 500 uL 80% methanol was added to each well. After incubation for 20 minutes at −80°C, cells were scraped off the plate, vortexed vigorously, and centrifuged at maximum speed. 250 uL of the resulting supernatant was dried under vacuum. Dried metabolites were stored at −80°C prior to mass spectrometry analysis.

### Cell lysis and immunoblotting

Cells were lysed in buffer containing 50 mM Tris pH 7.4, 1% Nonidet P-40, 0.25% sodium deoxycholate, 1 mM EDTA, 150 mM NaCl, 1 mM dithiothreitol, 1 mM sodium orthovanadate, 20 mM sodium fluoride, 10 mM beta-glycerophosphate, 10 mM sodium pyrophosphate, 2 ug ml^-1^ aprotinin, 2 ug ml^-1^ leupeptin and 0.7 ug ml^-1^ pepstatin.

Western blot analysis was performed using standard protocols, and the following commercial antibodies were used as probes: ASNS (Proteintech 14681-1-AP, 1:1000), ATF4 (Cell Signaling 11815, 1:500), phospho-T389 S6 kinase (Cell Signaling 9234, 1:500), S6 kinase (Cell Signaling 2708, 1:1000), phospho-S235/235 S6 ribosomal protein (Cell Signaling 4858, 1:3000), S6 ribosomal protein (Cell Signaling 2217, 1:1000), phospho-S1859 CAD (Cell Signaling 70307, 1:500), CAD (Cell Signaling 11933, 1:1000), and a-tubulin (Sigma T6074, 1:10,000).

### T cell killing assay

E0771 engineered to stably express SIINFEKL (lentiviral transduction followed by SIINFEKL selection via the Blue Fluorescent Protein tag) were seeded at 500 cells/well in 96 well plates overnight. Metformin, MT Cell Viability Substrate, and NanoLuc enzyme were added to media in the presence or absence of 1000 OT1 T cells and with or without 0.1 mM asparagine. Viability measurements were read after 1 hour (for baseline) and after 24-96 hours with RealTime-Glo™ MT Cell Viability Assay (Promega).

### CRISPR-mediated ASNS knockout

Guide oligos targeting CTCCATATGTATCTCTACCC (clone 1) and ATTGTCATAGAGGGCGTGCA (clone 2) were cloned into pSpCas9(BB)-2A-Puro. Cells were transfected with 600 ng of plasmid in a 24-well plate. After 24 hours, cells were selected with 1 ug/ml purmomycin for 24h and then refreshed with puromycin-free DMEM supplemented with 0.1 mM asparagine (DMEM+N). Cells were clonally selected in DMEM+N by plating 1 cell/well in 96-well plates and expanding following colony development. For ASNS restoration in ASNS KO cells, ASNS was expressed using a modified pCCL lentiviral vector under the CMV promoter.

### Mice

Mice were housed in pathogen-free facilities at University of California Los Angeles (UCLA). All animal experiments were approved by the UCLA Animal Research Committee (ARC), and we complied with all relevant ethical regulations while conducting animal experiments. We determined the number of mice required as the minimal number of mice necessary for reliable, significant results. Both male and female mice were used and no preference in mouse sex was given except when female mice were used for SUM159PT breast cancer xenografts.

### A549 and SUM159PT mouse tumour xenografts

5×10^6^ A549 or SUM159PT cells were injected subcutaneously into the flans of 7-9-week-old NSG mice. The mice were randomized, and treatments began when tumours reached an average of 200 mm^3^. For asparaginase experiments, pegylated-asparaginase (Jazz Pharmaceuticals, 5 IU/kg in 0.9% NaCl, total volume 100 ul) and 0.9% NaCl control were delivered by tail vein injections every 10 days. For asparagine-adjusted diets, mice were given an asparagine deficient diet (0% asparagine), a control diet (0.6% asparagine) or an asparagine-rich diet (4% asparagine). All diets were isonitrogenous and contained similar calorie densities. The diets were changed weekly.

Metformin-treated mice received 250 mg/kg/day metformin dissolved in water with 0.5% Splenda ad libitum in their drinking water, and the water was changed every 2-3 days. Control mice received 0.5% w/v Splenda in their drinking water. For IACS-010759 treatments, we resuspended 1 mg/kg IACS-010759 in 0.5% methylcellulose and delivered daily via oral gavage. Control mice were gavaged with 0.5% methylcellulose. Tumour volumes were measured every two days with calipers (using the formula mm^3^ = (LxW^2^)/2). Tumours were harvested when the first mouse reached our designated endpoint (1500 mm^3^), and snap frozen in liquid nitrogen. Serum was also collected and stored at −80 °C.

### Tumour metabolite extraction

Tumours were weighed and metabolites extracted in 1 ml 80% MeOH (−80 °C) using a Fisherbrand™ Bead Mill Homogenizer. Samples were spun twice at >17,000 *g* (4 °C) to remove debris. Extraction volumes equivalent to 3 mg of tissue were normalized to 500 ul with 80% MeOH and evaporated using a Genevac EZ-2 evaporator. Evaporated samples were stored at −80 °C.

### Serum metabolite extraction

20 ul of serum was mixed with 80 ul 100% MeOH (−80 °C) to give a final extraction concentration of 80% MeOH. Samples were centrifuged for 10 minutes at >17,000 *g* (4 °C) and 50 ul of each sample evaporated using a Genevac EZ-2 evaporator. Evaporated samples were stored at −80 °C.

### Tumour protein extraction

Proteins were extracted in 1 ml cell lysis buffer (see above) using a Fisherbrand™ Bead Mill Homogenizer. Samples were spun twice at >17,000 *g* (4 °C) to remove debris, and stored at −80 °C.

### KrasG12D;Lkb1-/-;Luc mice survival analysis

We used the Lox-Stop-Lox KrasG12D, Lkb1 lox/lox, Rosa26-Lox-Stop-Lox-Luciferase (KL) genetically engineered mouse model of lung cancer. Both male and female mice were used. We induced lung tumours by intranasal administration of 2.5 x 10^6^ plaque forming units of Adenoviral Cre (Gene Transfer Vector Core, University of Iowa) when mice were between 7-16 weeks of age. Treatment commenced two weeks after administration of Adenoviral Cre, when mice were randomized into two groups. One group was treated with a combination of food that contained 0% asparagine, and drinking water that had 250 mg/kg/day metformin dissolved in it (prepared as above). Control mice were fed food with 0.6% asparagine, and water with Splenda dissolved in it (as above). Treatment was continued until the mice reached a humane endpoint.

### Orthotopic Pancreatic Tumour Model

5×10^4^ 7940B pancreatic ductal carcinoma cells, derived from a C57BL/6J Kras^LSL-G12D/+^;Trp53^R172H/+^;Pdx1-Cre murine tumour, were injected into the pancreas tail of female syngeneic C57BL/6J mice in 50ul of a 2-1 matrigel:DMEM suspension. Tumour were allowed to establish for 8 days, then mice were randomized on to one of four treatment groups: 5% on sucrose water and treated every 3 days with PBS vehicle, 1.7mg/mL phenformin in 5% sucrose treated every 3 days with PBS vehicle, 2UI of Pegylated L-asparaginase delivered IP every 3 days on 5% sucrose water, or 2UI of Pegylated L-asparaginase delivered IP every 3 days on1.7mg/mL Phenformin in 5% sucrose water. Mice were sacrificed at a human endpoint after 10 days of treatment, then tumours were excised and weighed. Mice were maintained in specific pathogen free housing at constant ambient temperature and a 12-hour light cycle. Mice were used for orthotopic pancreas transplantations at 8-10 weeks of age. These experiments were conducted in accordance with the Office of Laboratory Animal Welfare and approved by the Institutional Animal Care and Use Committees of the University of Michigan.

### E0771 syngenic breast tumour growth and survival study

All mouse experiments were approved by the Cedars Sinai Medical Center Institutional Animal Care and Use Committee. Female 6-8-week-old C57BL/6J mice were purchased from JAX. All orthotopic injections had 2 x 10^5^ mouse mammary cancer cells resuspended in 50 ul of a 1:1 mix of PBS and Matrigel Basement Membrane (Corning). Primary tumour volume was measured using the formula mm^3^ = (LxW^2^)/2), in which *L* is length and *W* is width of the primary tumour. Mice were sacrified at our designated endpoint (1500 mm^3^). Metformin was dissolved in drinking water (250 mg/kg/day) supplemented with 0.5 g Splenda in 100 ml water for palatability (0.5% w/v Splenda). Splenda and Metformin was administered with drinking water and changed every 2-3 days. For L-asparagine-adjusted diets, mice were given an asparagine deficient diet (0% asparagine), or an asparagine-rich diet (4% asparagine). All diets were isonitrogenous and contained similar calorie densities. Mice were intraperitoneally injected with anti-mouse PD1(CD279) and rat IgG2a isotype control (InVivoMAb) at 200 μg/injection once primary tumour reached 7mm at any one dimension. Antibody injections were given every 3 days for up to 5 injections then once a week until experimental completion. Cyclophosphamide monohydrate were injected retro-orbitally at 30 mg/kg once the primary tumour reached 10mm at any one dimension (Thermo Fisher Scientific). Weekly injections of Cyclophosphamide occurred until experimental completion. Animals were assigned to treatment groups randomly.

### Mass spectrometry-based metabolomics analysis

Dried metabolites were reconstituted in 100 µL of a 50% acetonitrile(ACN) 50% dH20 solution. Samples were vortexed and spun down for 10 min at 17,000g. 70 µL of the supernatant was then transferred to HPLC glass vials. 10 µL of these metabolite solutions were injected per analysis. Samples were run on a Vanquish (Thermo Scientific) UHPLC system with mobile phase A (20mM ammonium carbonate, pH 9.7) and mobile phase B (100% ACN) at a flow rate of 150 µL/min on a SeQuant ZIC-pHILIC Polymeric column (2.1 × 150 mm 5 μm, EMD Millipore) at 35°C. Separation was achieved with a linear gradient from 20% A to 80% A in 20 min followed by a linear gradient from 80% A to 20% A from 20 min to 20.5 min. 20% A was then held from 20.5 min to 28 min. The UHPLC was coupled to a Q-Exactive (Thermo Scientific) mass analyzer running in polarity switching mode with spray-voltage=3.2kV, sheath-gas=40, aux-gas=15, sweep-gas=1, aux-gas-temp=350°C, and capillary-temp=275°C. For both polarities mass scan settings were kept at full-scan-range=(70-1000), ms1-resolution=70,000, max-injection-time=250ms, and AGC-target=1E6. MS2 data was also collected from the top three most abundant singly-charged ions in each scan with normalized-collision-energy=35. Each of the resulting “.RAW” files was then centroided and converted into two “.mzXML” files (one for positive scans and one for negative scans) using msconvert from ProteoWizard^28^. These “.mzXML” files were imported into the MZmine 2 software package^29^. Ion chromatograms were generated from MS1 spectra via the built-in Automated Data Analysis Pipeline (ADAP)^30^ chromatogram module and peaks were detected via the ADAP wavelets algorithm. Peaks were aligned across all samples via the Random sample consensus aligner module, gap-filled, and assigned identities using an exact mass MS1(+/-15ppm) and retention time RT (+/-0.5min) search of our in-house MS1-RT database. Peak boundaries and identifications were then further refined by manual curation. Peaks were quantified by area under the curve integration and exported as CSV files. If stable isotope tracing was used in the experiment, the peak areas were additionally processed via the R package AccuCor^31^ to correct for natural isotope abundance. Peak areas for each sample were normalized by the measured area of the internal standard trifluoromethanesulfonate (present in the extraction buffer) and by the number of cells present in the extracted well.

## Data Availability

The authors declare that the data supporting the findings of this study are available within the paper.

---

**Supplementary Figure 1.**
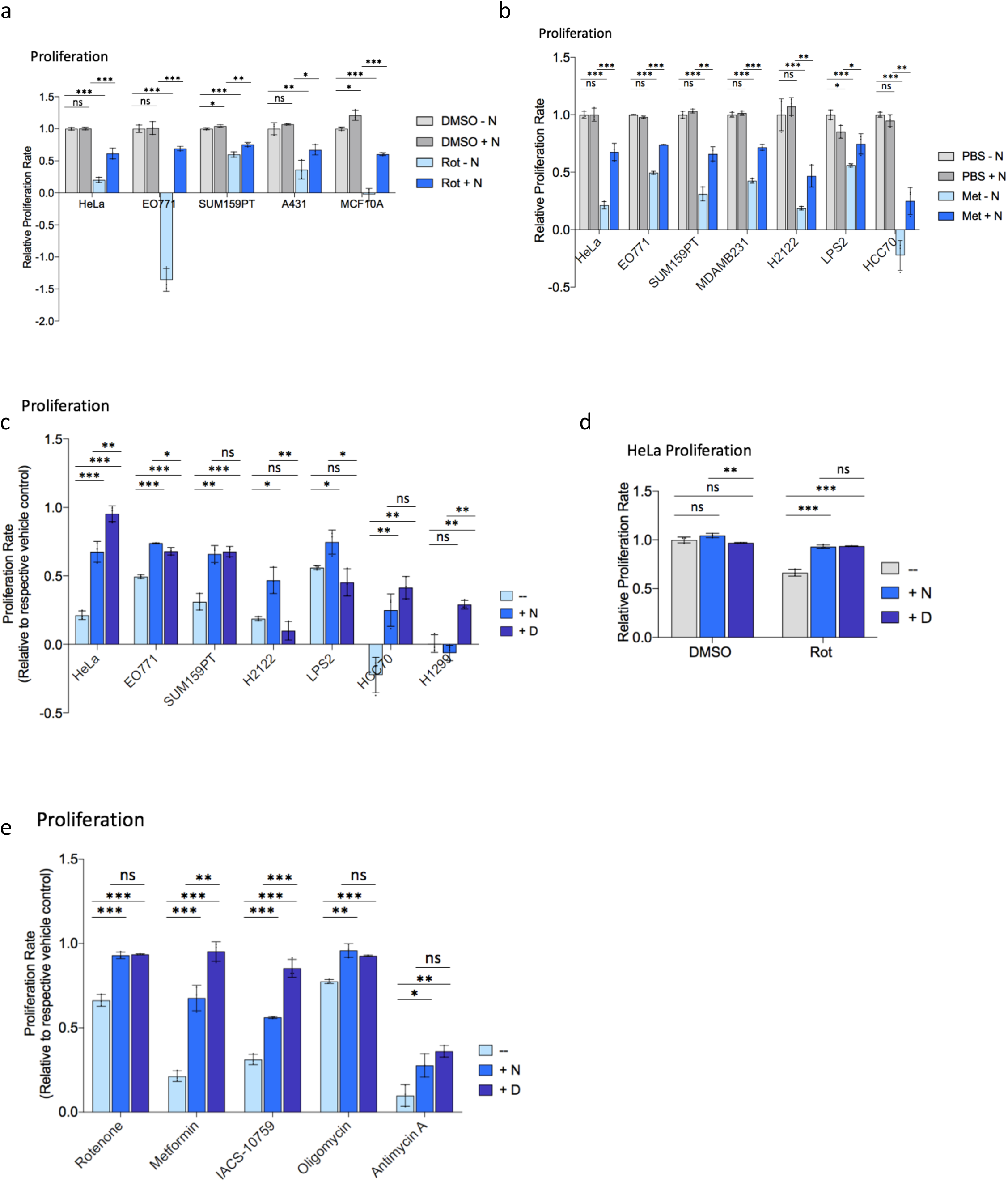
Either asparagine or aspartate enables proliferation in the context of ETC inhibition. **a**, Relative proliferation rate of indicated cell lines with rotenone (Rot) or DMSO treatment in the presence or absence of 0.1 mM exogenous asparagine (N). **b**, Relative proliferation rate of indicated cell lines with metformin (Met) or vehicle control (PBS) treatment in the presence or absence of 0.1 mM exogenous asparagine (N). **c,** Proliferation rate of indicated cell lines with metformin treatment in the presence or absence of 0.1 mM exogenous asparagine (N) or 20 mM aspartate, relative to respective PBS control proliferation in unsupplemented DMEM. **d**, Relative HeLa cell proliferation rate with rotenone or DMSO treatment in the presence or absence of 0.1 mM exogenous asparagine (N) or 20 mM aspartate (D). **e**, Relative HeLa cell proliferation rate with the indicated ETC inhibitor in the presence or absence of 0.1 mM exogenous asparagine (N) or 20 mM aspartate (D). Data are mean +/- s.d. (n = 3 independent experiments). P value determined by unpaired two-tailed t-test: *p<0.05; **p<0.01; ***p<0.001; ns, not significant.

**Supplementary Figure 2.**
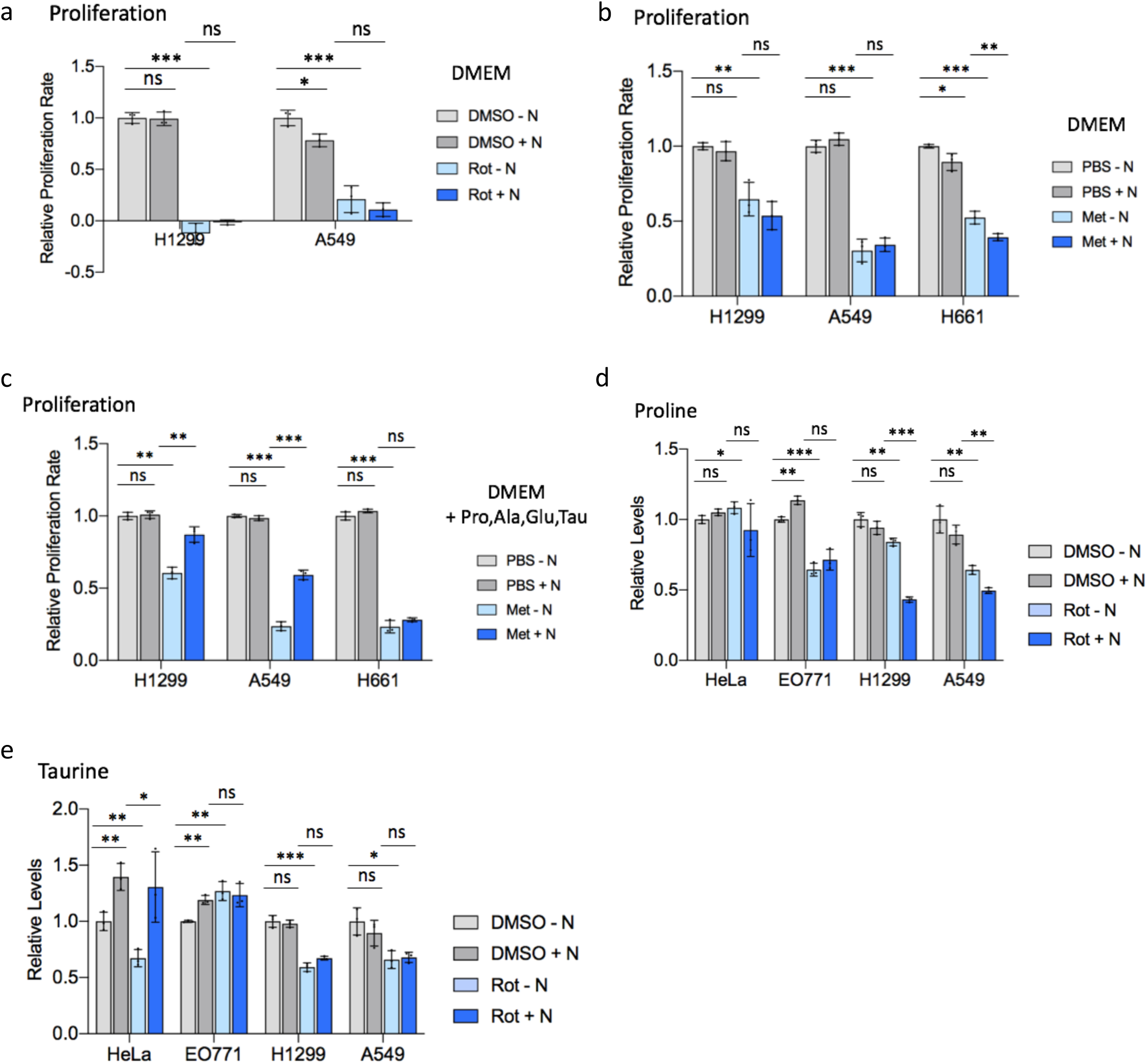
Cell environment can affect the ability of asparagine to restore proliferation in the context of ETC inhibition. **a**, Relative proliferation rate of indicated cell lines with rotenone (Rot) or DMSO treatment in standard pyruvate-free DMEM in the presence or absence of 0.1 mM exogenous asparagine (N). **b**, Relative proliferation rate of indicated cell lines with metformin (Met) or PBS treatment in standard pyruvate-free DMEM in the presence or absence of 0.1 mM exogenous asparagine (N). **c**, Relative proliferation rate of indicated cell lines with metformin or PBS treatment in pyruvate-free DMEM supplemented with 0.4 mM proline, 0.5 mM alanine, 0.1 mM glutamate, and 0.15 mM taurine in the presence or absence of 0.1 mM exogenous asparagine (N). Relative levels of intracellular proline (**d**) and taurine (**e**) in the indicated cell line 6 hours post-treatment with rotenone or DMSO in standard pyruvate-free DMEM in the presence or absence of 0.1 mM exogenous asparagine (N). Data are mean +/- s.d. (n = 3 independent experiments). P value determined by unpaired two-tailed t-test: *p<0.05; **p<0.01; ***p<0.001; ns, not significant.

**Supplementary Figure 3.**
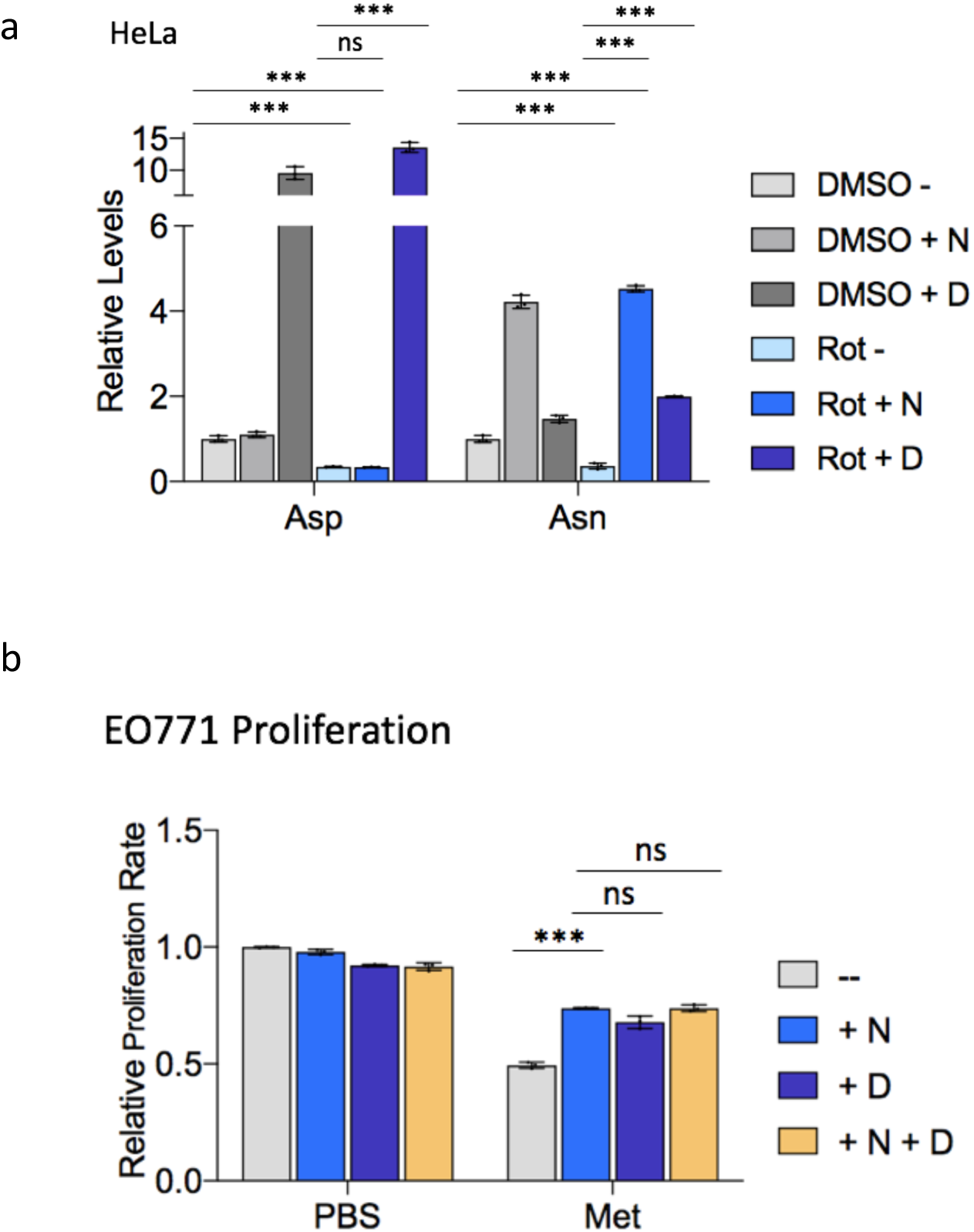
Asparagine and aspartate may rescue proliferation through a common mechanism. **a**, Relative levels of intracellular asparagine (Asn) and aspartate (Asp) in HeLa cells 48 hours post-treatment with 50 nM rotenone (Rot) or DMSO presence or absence of 0.1 mM exogenous asparagine (N) or 20 mM aspartate (D). **b,** Relative E0771 cell proliferation rate with 5 mM metformin (Met) or PBS treatment in the presence or absence of 0.1 mM exogenous asparagine (N), 20 mM aspartate (D), or a combination of 0.1 mM asparagine and 20 mM aspartate. Data are mean +/- s.d. (n = 3 independent experiments). P value determined by unpaired two-tailed t-test: *p<0.05; **p<0.01; ***p<0.001; ns, not significant.

**Supplementary Figure 4.**
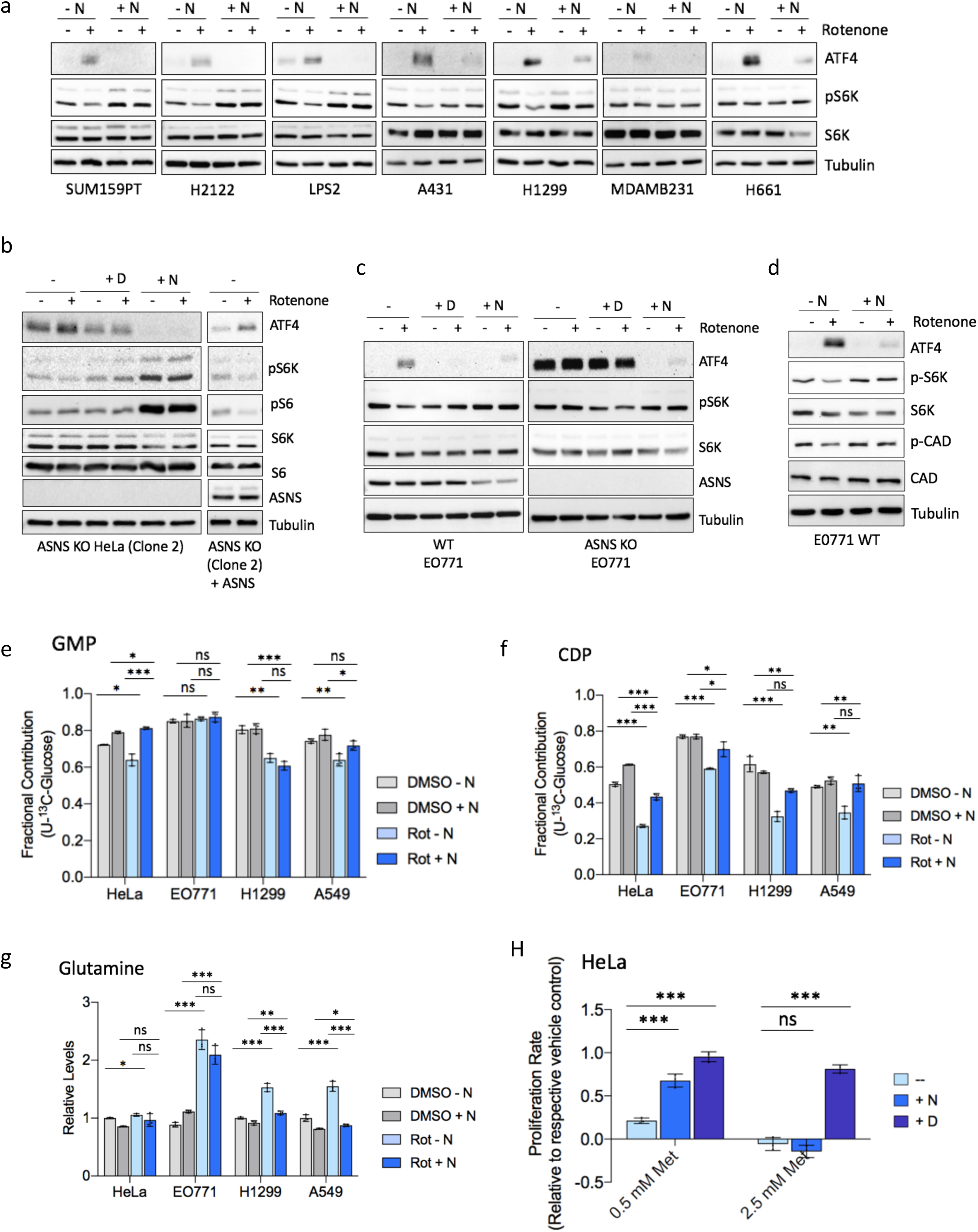
Asparagine restores ATF4 and mTORC1 activities in the context of ETC inhibition. **a**, Immunoblot of lysates after 6 hours post-rotenone (rot) (50 nM) or DMSO treatment. Lysates were immunoblotted for ATF4, mTORC1 activation markers phospho-Thr389 S6K, total S6K, and tubulin. **b**, Left, Immunoblot of HeLa ASNS KO (clone 2; see Fig. 1c for ASNS KO clone 1) lysates 6 hours post-treatment with 50 nM rotenone or DMSO in the presence or absence of 20 mM aspartate (D) or 0.1 mM asparagine (N); Right, ASNS was restored in HeLa ASNS KO cells with CMV-driven ectopic expression. Immunoblot shows lysates 6 hours post-treatment with 50 nM rotenone or DMSO in unsupplemented DMEM. **c**, Immunoblot of E0771 WT or ASNS KO lysates 3 hours post-treatment with 50 nM rotenone or DMSO in the presence or absence of 0.1 mM asparagine (N) or 20 mM aspartate (D). **d**, Immunoblot of E0771 WT lysates 3 hours post-treatment with 50 nM rotenone or DMSO in the presence or absence of 0.1 mM asparagine (N). Lysates were immunoblotted for ATF4, mTORC1 activation markers phospho-Thr389 S6K and phospho-Ser1859 CAD, total S6K and CAD, and tubulin. Fractional contribution of U-^13^C-glucose to GTP (**e**) and CDP (**f**) in the indicated cell line 6 hours post-treatment with rotenone or DMSO presence or absence of 0.1 mM exogenous asparagine. Medium was replaced with DMEM containing 10 mM U-^13^C-glucose at the same time as rotenone treatment. **g**, Relative levels of intracellular glutamine in the indicated cell line 6 hours post-treatment with rotenone or DMSO presence or absence of 0.1 mM exogenous asparagine (N). **h**, HeLa cell proliferation rate with either 0.5 or 2.5 mM metformin (Met) treatment in the presence or absence of 0.1 mM exogenous asparagine (N) or 20 mM aspartate, relative to PBS control proliferation in unsupplemented DMEM. For **e-h**, data are mean +/- s.d. (n = 3 independent experiments). P value determined by unpaired two-tailed t-test: *p<0.05; **p<0.01; ***p<0.001; ns, not significant.

**Supplementary Figure 5.**
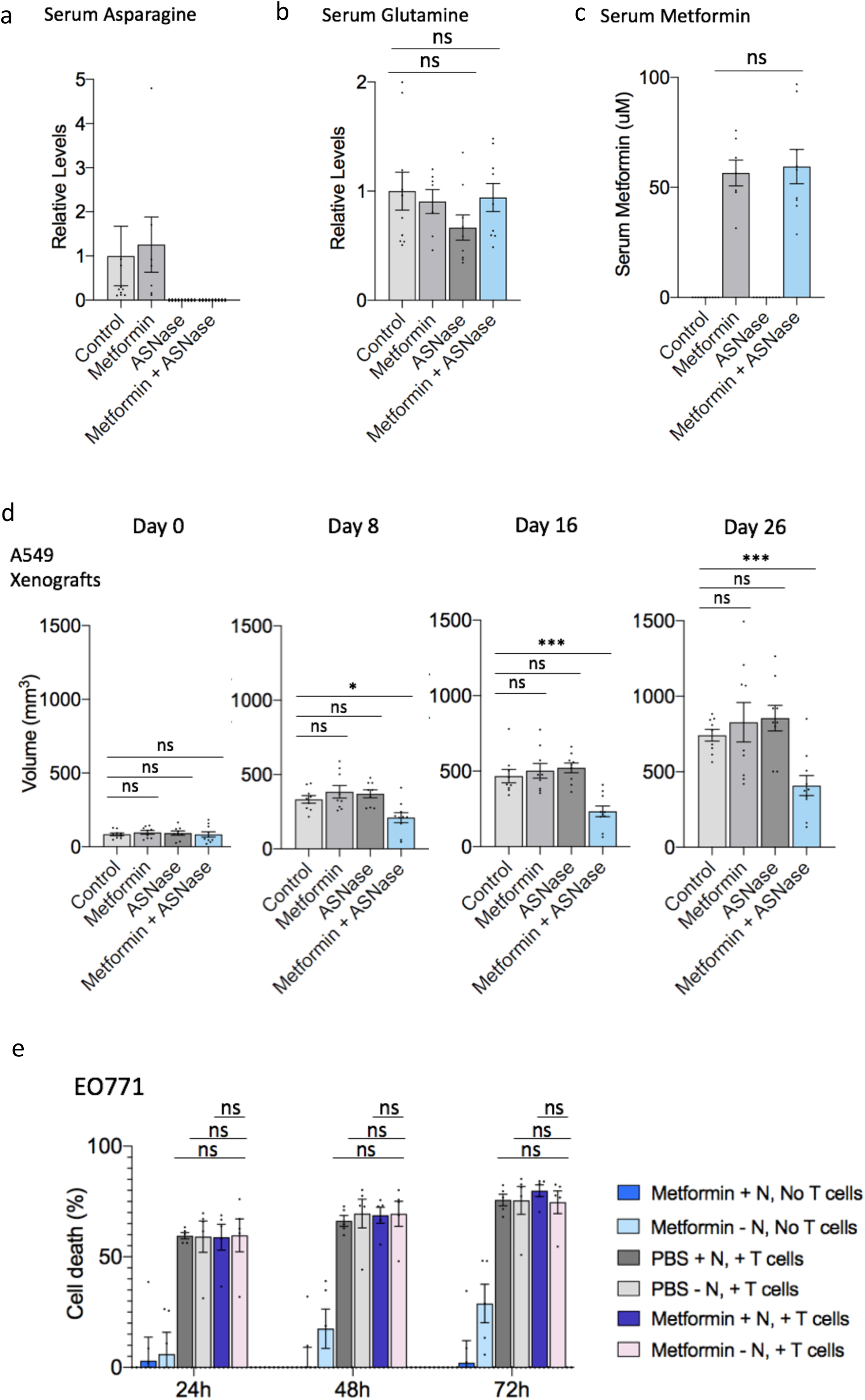
Combining metformin with asparaginase inhibits tumour growth. Serum asparagine (**a**), glutamine (**b**) and metformin (**c**) levels from mice treated with metformin, asparaginase (ASNase), the combination, or vehicle controls. n = 7-10. **d**, Tumour volume (mm^3^) of A549 subcutaneous tumour xenografts at days 0, 8, 16, and 26 post-initiation of treatment with metformin (250 mg/kg/day), asparaginase (ASNase) (5 IU/kg), the combination, or vehicle controls as determined by caliper measurements. n = 9-10. **e**, Cell death of E0771 cells with metformin treatment in the presence of absence of 0.1 mM exogenous asparagine (N) and in the presence (grey and purple bars) or absence (blue bars) of OT1 T cells, n = 5. Data are mean +/- s.e.; P value determined by unpaired two-tailed t-test: *p<0.05; **p<0.01; ***p<0.001; ns, not significant.

**Supplementary Figure 6.**
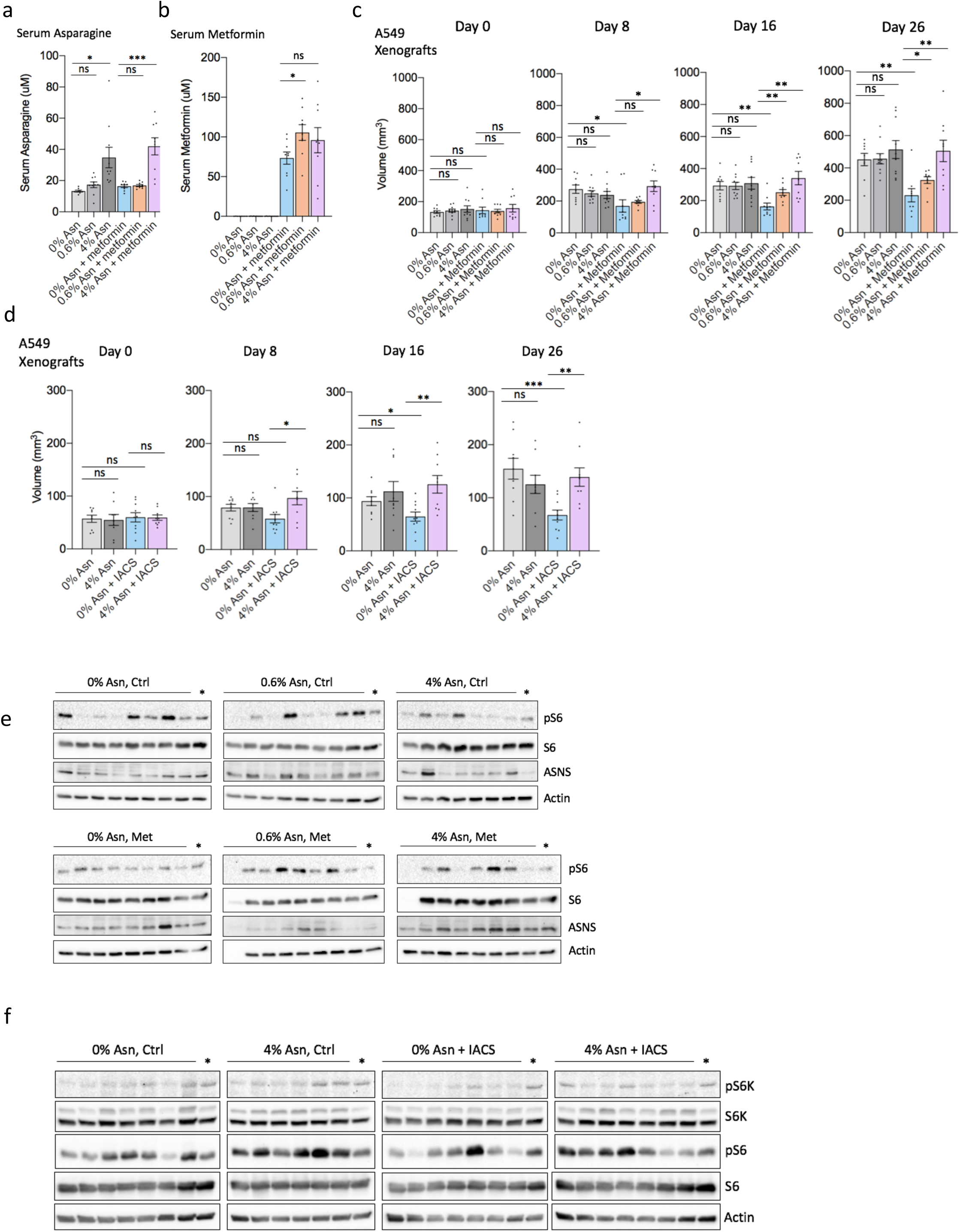
Combining metformin with dietary asparagine restriction inhibits tumour growth. Serum asparagine **(a)** and metformin **(b)** levels from mice treated with or without metformin and diets containing 0%, 0.6%, or 4% asparagine (Asn). **c**, Tumour volume (mm^3^) of A549 subcutaneous tumour xenografts at days 0, 8, 16, and 26 post-initiation of treatment with or without metformin and diets containing 0%, 0.6%, or 4% asparagine (Asn). **d**, Tumour volume (mm^3^) of A549 subcutaneous tumour xenografts at days 0, 8, 16, and 26 post-initiation of treatment with or without IACS-010759 (IACS) and diets containing 0% or 4% asparagine (Asn). **e**, Immunoblot of A549 tumour lysates from mice treated with or without metformin and fed a diet containing 0%, 0.6%, or 4% asparagine. Each lane contained lysate from individual tumours within each treatment group. Lysates were immunoblotted for mTORC1 activation marker phospho-Ser235/6 S6, total S6, ASNS, and actin. The asterisk (*) indicates a reference lysate (taken from the largest of the “4% Asn, Ctrl” group) that was run on each gel. **f**, Immunoblot of A549 tumour lysates from mice treated with or without IACS-010759 (IACS) and fed a diet containing 0% or 4% asparagine. Each lane contained lysate from individual tumours within each treatment group. Lysates were immunoblotted for mTORC1 activation marker phospho-Ser235/6 S6, total S6, ASNS, and actin. The asterisk (*) indicates a reference lysate (taken from the largest of the “4% Asn, Ctrl” group) that was run on each gel. For **a-d**, data are mean +/- s.e.; P value determined by unpaired two-tailed t-test: *p<0.05; **p<0.01; ***p<0.001; ns, not significant.

**Supplementary Figure 7.**
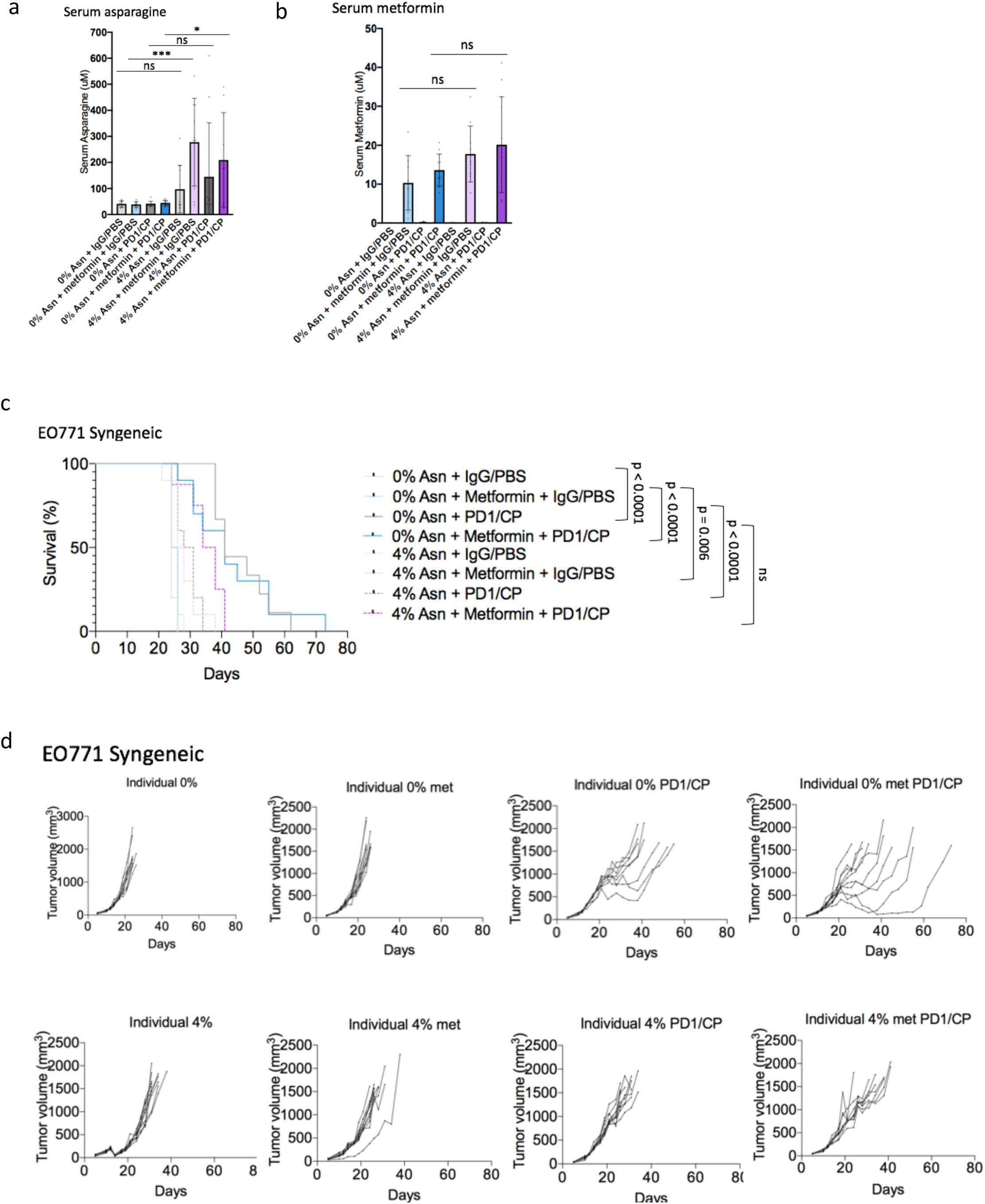
Dietary asparagine restriction synergizes with PD1 inhibition and chemotherapy. Endpoint serum asparagine **(a**) and metformin **(b)** from mice harboring syngeneic E0771 mammary tumours and treated with or without metformin and/or a combination of inhibitory PD1 antibody and cyclophosphamide (CP) and fed a diet containing 0% or 4% asparagine (Asn), n = 10. **c**, Kaplan-Meier survival curve indicating percentage of mice with syngeneic E0771 mammary tumours surviving with the indicated treatments, n = 10. **d**, Individual tumour growth curves of E0771 tumours with the indicated treatment. For **a-b**, data are mean +/- s.e.; P value determined by unpaired two-tailed t-test. For **c**, P-value was calculated by Mantel-Cox test. *p<0.05; **p<0.01; ***p<0.001; ns, not significant.

**Supplementary Figure 8.**
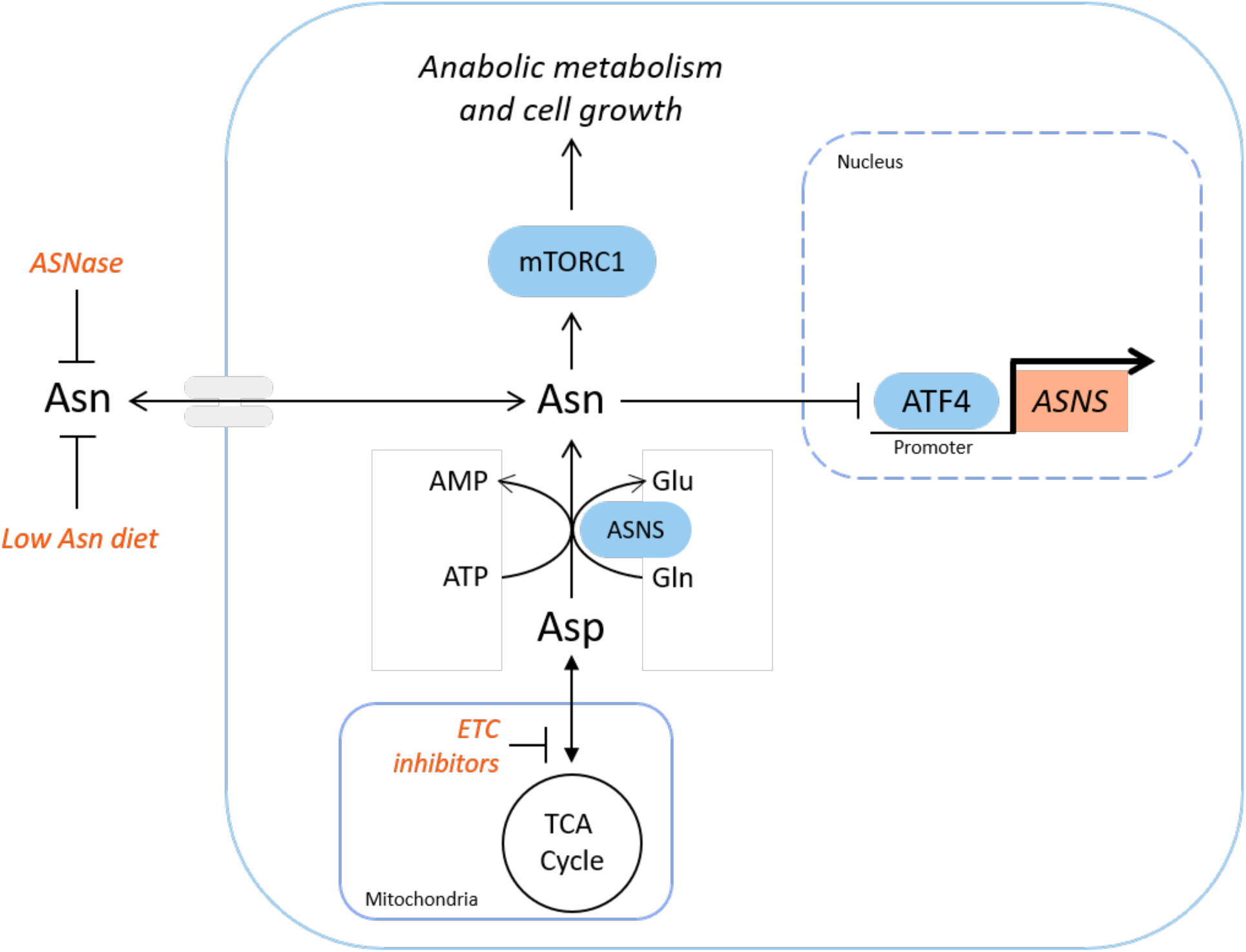
Asparagine signals mitochondrial respiration to mTORC1 and ATF4 and can be targeted to impair tumor growth. Schematic showing communication of mitochondrial respiration to mTORC1 and ATF4 by aspartate-derived asparagine. Tumor growth can be impaired by combining ETC inhibition, which blocks *de novo* asparagine synthesis, with either asparaginase or dietary asparagine restriction, which limit asparagine consumption.

**Supplementary Table I:**
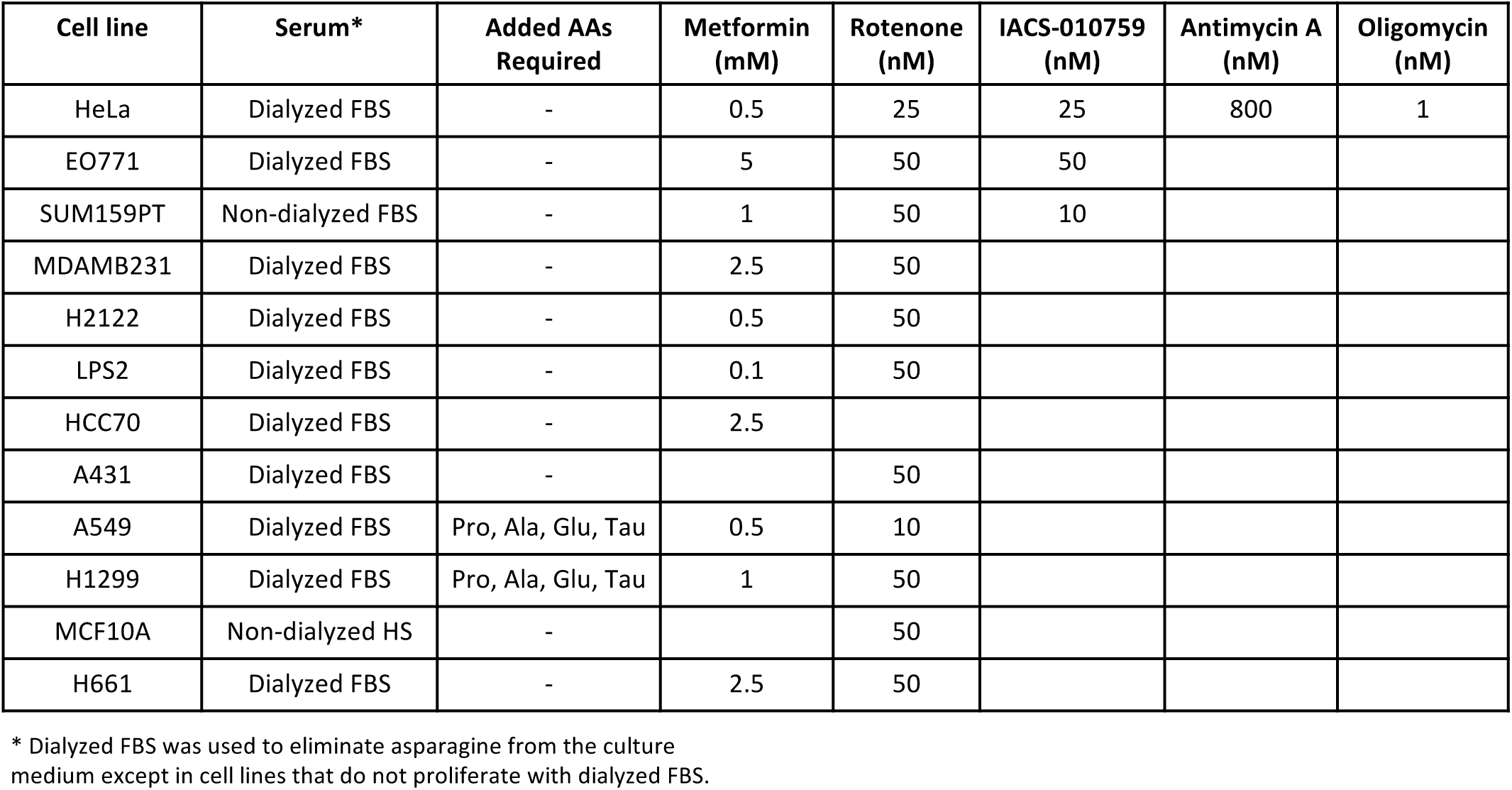
Cell line-specific conditions for ETC inhibition

| Cell line | Serum* | Added AAs Required | Metformin (mM) | Rotenone (nM) | IACS-010759 (nM) | Antimycin A (nM) | Oligomycin (nM) |
| --- | --- | --- | --- | --- | --- | --- | --- |
| HeLa | Dialyzed FBS | - | 0.5 | 25 | 25 | 800 | 1 |
| EO771 | Dialyzed FBS | - | 5 | 50 | 50 |  |  |
| SUM159PT | Non-dialyzed FBS | - | 1 | 50 | 10 |  |  |
| MDAMB231 | Dialyzed FBS | - | 2.5 | 50 |  |  |  |
| H2122 | Dialyzed FBS | - | 0.5 | 50 |  |  |  |
| LPS2 | Dialyzed FBS | - | 0.1 | 50 |  |  |  |
| HCC70 | Dialyzed FBS | - | 2.5 |  |  |  |  |
| A431 | Dialyzed FBS | - |  | 50 |  |  |  |
| A549 | Dialyzed FBS | Pro, Ala, Glu, Tau | 0.5 | 10 |  |  |  |
| H1299 | Dialyzed FBS | Pro, Ala, Glu, Tau | 1 | 50 |  |  |  |
| MCF10A | Non-dialyzed HS | - |  | 50 |  |  |  |
| H661 | Dialyzed FBS | - | 2.5 | 50 |  |  |  |

\* Dialyzed FBS was used to eliminate asparagine from the culture medium except in cell lines that do not proliferate with dialyzed FBS.

